# Strain intrinsic properties and environmental constraints together shape *Escherichia coli* dynamics and diversity over a twenty-year human gut time series

**DOI:** 10.1101/2024.02.21.581337

**Authors:** Bénédicte Condamine, Thibaut Morel-Journel, Florian Tesson, Guilhem Royer, Mélanie Magnan, Aude Bernheim, Erick Denamur, François Blanquart, Olivier Clermont

## Abstract

*Escherichia coli* is an increasingly antibiotic-resistant opportunistic pathogen. Few data are available on its ecological and evolutionary dynamics in its primary commensal niche, the vertebrate gut. Using Illumina and/or Nanopore technologies, we sequenced whole genomes of 210 *E. coli* isolates from 22 stools sampled during a 20-year period from a healthy man (ED) living in Paris, France. All phylogroups, except C, were represented, with a predominance of B2 (34.3%), followed by A and F (19% each) phylogroups. Thirty-five clones were identified based on their haplogroup and pairwise genomic single nucleotide polymorphism distance and classified in three phenotypes according to their abundance and residence time: 25 sub-dominant/transient (52 isolates), five dominant/transient (48 isolates) and five dominant/resident (110 isolates). Four over five dominant/resident clones belonged to B2 and closely related F phylogroups, whereas sub-dominant/transient clones belonged mainly to B1, A and D phylogroups. The long residence times of B2 clones seemed to be counterbalanced by lower colonization abilities. Clones with larger within-host frequency persisted for longer in the host. By comparing ED strain genomes to a collection of commensal *E. coli* genomes from 359 French individuals, we identified ED-specific genomic properties including a set of genes involved in a metabolic pathway (*mhp* cluster) and a very rare antiviral defense island. The *E. coli* colonization within the gut microbiota was shaped by both the intrinsic properties of the strain lineages, in particular longer residence of phylogroup B2, and the environmental constraints such as diet or phages.

## Introduction

*Escherichia coli* intestinal (diarrhea) (1) and extra-intestinal (urinary tract infection, septicemia, pneumonia, deep suppuration) (2) infections represent a major public health problem due to the frequency and the severity of these diseases. *E. coli* is also becoming increasingly antibiotic resistant (3). However, the normal niche of *E. coli* is the gut of vertebrates, where it lives as a commensal (4). This species is thus acting mostly as an opportunistic pathogen. Selection for antimicrobial resistance happens for a large part in the gut, as a commensal (5). Strikingly, despite the urgent need to understand the normal ecology of *E. coli* to fight or manage virulence and resistance, little knowledge is available on its normal lifestyle and especially on the *E. coli* population changes over time (6), as compared to the abundant literature on *E. coli* infections. Whole genome sequences of well-documented dominant commensal isolates are still rare compared to clinical ones.

*E. coli* population structure and dynamics in the gut of healthy individuals has been addressed mostly in humans since the beginning of the 20^th^ century, using in each epoch the available technologies: serotyping (7–11), multilocus enzyme electrophoresis (12,13), PCR-based approaches, pulsed-field gel electrophoresis, multilocus sequence typing, ribotyping, amplicon sequencing of the flagellin gene and shotgun metagenomics (6,14–19). Most of these studies were performed on one to 10 subjects over a few months (maximum 4 years (10)) studying in general 10 colonies per sample. Some consensus emerged from these studies: (i) *E. coli* can present two extreme phenotypes associated with colonization ability: those persisting for short periods (days to weeks), the “transients” and those persisting for longer periods (but not on very long timescales), the “residents” (20), (ii) a single “dominant” strain (90 to 100% within sample) is present in the majority of the samples whereas in other samples several “minor” or “sub-dominant” strains are present, (iii) the resident strains correspond to specific phylogenetic lineages often involved in extra-intestinal infections belonging to phylogroups B2 (sequence types (STs) 73, 95, 131) and F (ST59) and (iv) there is an important inter-individual variability. Similar results were obtained in non-human mammals, either in the wild in brushtail possums in Australia (21) or in cattle in Zimbabwean cows (22).

More recently, whole genome sequencing (WGS) of single isolates was used to study the evolution of a clone between the members of a household over three years (23) and we have previously studied the *E. coli* carriage of a healthy man (ED) living in Paris, France, during one year (24). Both studies found an observed mutation rate per base per year of about 2×10^-7^, in the range of what was observed in the *in vitro* long-term experiment of Richard Lenski (25). However, at the opposite of the Lenski experiment that shows strong selection (26), *in vivo* evolution appears to be dominated by neutral changes.

In the present work, we extended the study of ED *E. coli* carriage to 20 years (2001–2020). We analyzed 22 stool samples irregularly spaced over these 20 years, including the three first samples already published (24). For each new sample, we whole-genome sequenced 10 *E. coli* colonies using Illumina and/or Nanopore technologies. In addition, we compared the ED strain collection to an epidemiologically relevant collection of 359 *E. coli* isolates from the stools of 359 healthy individuals living in France and sampled between 2000 and 2017 to search for ED individual strain-specific traits.

## Materials/subjects and methods

### Ethics

The study was approved by the ethics evaluation committee of Institut National de la Santé et de la Recherche Médicale (INSERM) (CCTIRS no. 09.243, CNIL no. 909277, and CQI no. 01-014). The authors were given consent by the subject.

### Strain samples

During a 20-year period (2001–2020), we irregularly sampled the stools of a healthy man (ED) (44-64 year-old) living in Paris (see supplementary online material (SOM) for more details on ED), with in 2019 monthly sampling during seven months and weekly sampling during one month, nested in the monthly collect (Fig. 1 and Table S1). Ten isolates appearing as *E. coli* were randomly selected after plating the feces on Drigalski medium without attempting to maximize phenotypic diversity (see SOM) and stored at – 80°C in glycerol (220 isolates for 22 samples). The isolates of the three first samplings have been published previously (24).

**Fig. 1.**
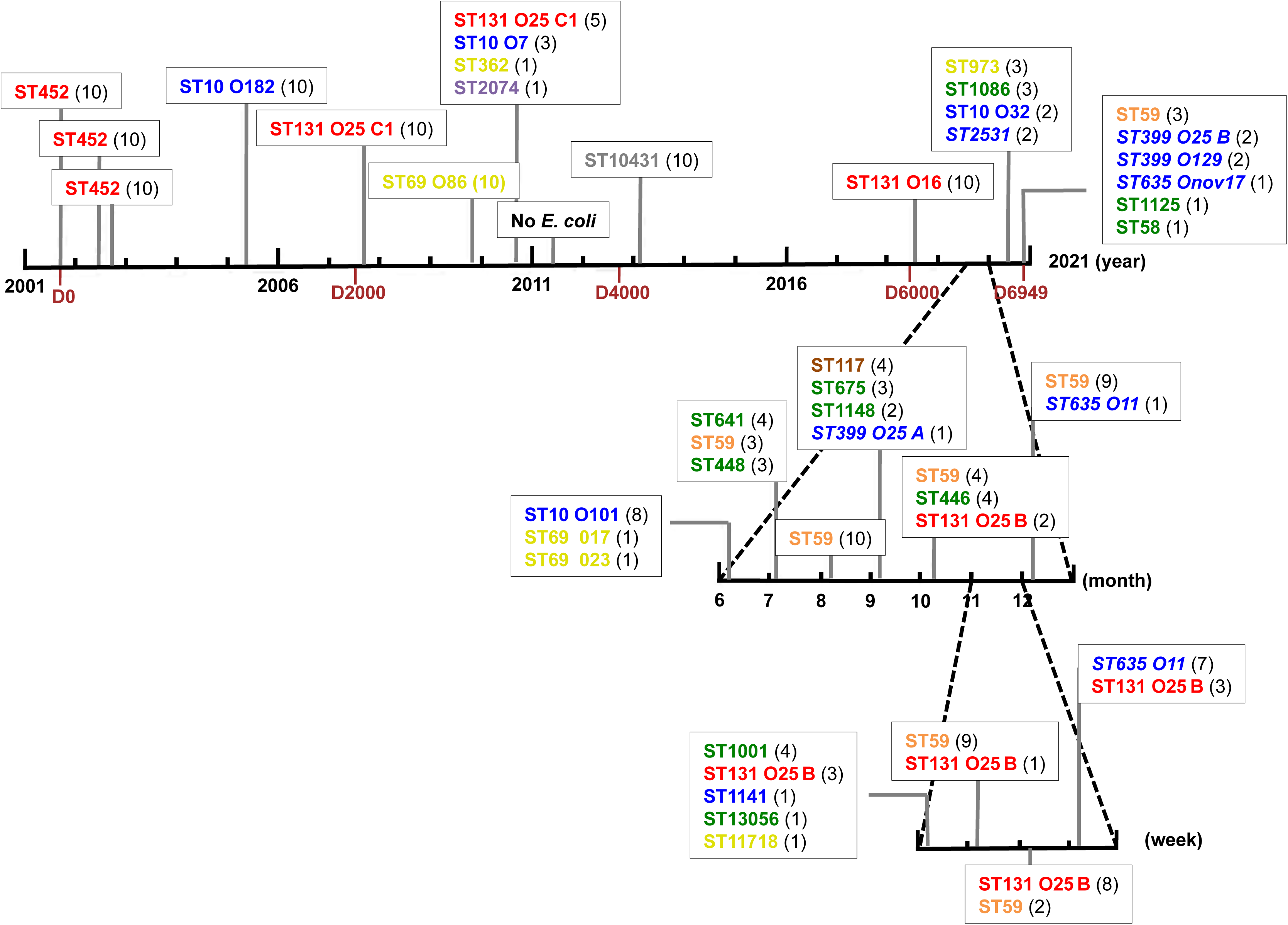
Global representation of the study with the dates of sampling and the identified clones of the 22 stool samples performed during a 20 year-period. Ticks on the main (top) horizontal line correspond to one year intervals. Two periods with more frequent sampling are represented as sublines nested on the main line, with ticks defining monthly (middle line) and weekly (bottom line) intervals. Among the 22 samples, only one in 2011, did not yield any *E. coli* but a unique clone of *Hafnia alvei*. For the sake of clarity the *E. coli* clones are identified in a simplified way by their STs, and when necessary, their O-groups and their sub-types. The colors correspond to their phylogroup (A=blue, B1=green, B2=red, D=yellow, E=violet, F=orange, G=brown, H=grey). The number of isolates belonging to the clone is indicated in brackets. The first sample corresponds to day 0 (D0) in 2001 and the last one to day 6949 (D6949) in 2020. The clones bearing the dGTPase and/or Detocs defense system(s) are indicated in italics.

### Genome sequencing

All ED isolates were sequenced using Illumina technology (short reads) on MiniSeq sequencer. The genomes were assembled using SPAdes v3.13.1 and standard parameters, and then annotated with Prokka 1.14.5 (27,28). Sequences have been deposited at the ENA under the bioproject PRJEB45090 (Table S1). One strain from the first sample, ED1a, has also been sequenced using standard Sanger technology and assembled in two single molecules, the chromosome and the plasmid (29). This sequence was then used in all the genomic analysis. Nanopore sequencing (long reads) was performed on one selected isolate per clone for 9 clones harboring at least 8 isolates. This sequenced isolate was selected randomly, in most of the cases within the first sample of the temporal series, and was considered as the nanopore reference isolate.

### Genomes used for comparison

Commensal *E. coli* strains were gathered from stools of 359 healthy adults living in the Paris area or Brittany (both locations in the Northern part of France) between 2000 to 2017. One strain per individual was sequenced as above (30). The 1,777 RefSeq *E. coli* genomes of high quality were downloaded from NCBI (July 2022).

### Genome analyses

Genome typing was performed as previously described (31) (https://github.com/iame-researchCenter/Petanc), including species identification, phylogrouping (32), multi-locus sequence type (MLST) determination according to the Warwick University and Pasteur Institute schemes, *in silico* serotyping (33,34) and *fimH* allele determination (35). The presence of virulence associated genes (coding for adhesins, iron capture systems, toxins, protectins) and resistance genes was determined as previously described with ABRicate 1.0.0, based on a custom database and Resfinder, respectively (36). The chromosomal or plasmidic location of a gene was determined using PlaScope 1.3.1 (37). We also searched for point mutations responsible for resistance using PointFinder 4.2.2 (38). The phenotypic resistance pattern of the strains to main classes of antibiotics was deduced from the presence of corresponding genes or mutations using the validated approach of (39).

Pan and core genomes were computed using PPanGGOLiN 1.2.105 with default parameters (40). Pairwise single nucleotide polymorphism (SNP) distance matrices from multiple sequence alignments were performed using pairsnp (https://github.com/gtonkinhill/pairsnp). Clones were defined according to the following strategy (41). First, isolates were grouped by their haplogroup, defined as a combination of their sequence types (ST) according to the Warwick University and Institute Pasteur schemes, their serotype (O:H), and their *fimH* allele. Then, the distribution of core genome pairwise SNP distances was determined to identify a cut-off where few/no values were observed, that can be used as a threshold to define clones. The SNP distances between isolates were compared with their haplogroup, allowing a data-based definition of the clone.

The genome-wide association studies (GWAS) were performed with Scoary 1.6.16 (42) (with default parameters). A maximum likelihood phylogenetic tree was inferred based on a core genome alignment using IQ-TREE 2.2.0 (43) and the GTR+F+I+G4 model (1000 bootstraps) rooted on NILS31 (*Escherichia* clade I). Visualization and annotation of the trees were performed with iTOL (44). Defense system detection was done using DefenseFinder models v1.2.3 (45). Additional methods to analyze the genetics of the defense island and *mhp* cluster are detailed in the SOM.

Illumina and nanopore contigs were assembled using Unicycler v0.4.9b (46) and annotated with Prokka 1.14.5 (28). We then filtered the mutations identified by breseq 0.33.2 (47) as in (24). When filtering the outputs of breseq, we looked at the predicted SNPs and short indels in the chromosomes only. We removed mutations when they were less than 51 bp apart. These clustered mutations are indeed usually caused by erroneously mapped reads.

### Bayesian inference of residence times

We inferred the posterior distribution of residence time for each observed clone, then grouping clones by sequence types and by phylogroups.

We assumed the residence time of a clone is exponentially distributed with a mean residence time T = 1/λ that we estimate. The exponential distribution is a simple, one-parameter distribution that emerges when the clearance rate is constant in time; we chose it for its simplicity in the absence of detailed data that could inform on potentially more complex distributions. To use prior information on residence times detailed by published studies, we adopted a Bayesian inference approach. For the prior distribution of the residence time we chose a log-normal distribution with parameters 3 and 1.5. This gives a mean of approximately 20 days and a 95% higher prior density interval of 1 to 400 days, in accordance with the overall mean and range of residence times observed in the longitudinal follow-up of 8 healthy volunteers (48).

For each episode of carriage of a clone, we defined the minimum and maximum residence times T_max_ and T_min_ from the date when it was first and last observed, and accounting for left and right censoring. The associated likelihood is the probability that the event of clearance took place in the time interval after colonization by the clone:

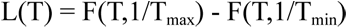

Where F(T,λ)=1-exp(-Tλ) is the cumulative distribution function of the exponential distribution.

For each clone we used the associated likelihood and the prior to define the posterior distribution of T, determine the mode of the posterior distribution of carriage time, as well as the 95% higher posterior density interval. We repeated the same analysis when grouping all clones of the same ST together assuming independence between carriage times of these distinct clones. We repeated the same analysis when grouping all clones of the same phylogroup together.

Many clones are observed only once. These clones have T_min_= 0 but do not have the same T_max_. However, in all cases the posterior mode of residence time was inferred to be at Ť=2.1 days.

### Analysis of the within-clone diversity

We studied the within-clone diversity, based on the mutations accumulated by each clone during its residence in the ED (see SOM for details on the method). On the one hand, we estimated an observed mutation rate based on differences between isolates of a given clone. On the other hand, we compared the observed differences to simulations of the Wright-Fisher model to assess the expected effective population size in ED under the hypothesis of neutrality.

## Results

Twenty-two stool samples from the ED subject, corresponding to 220 isolates (10 isolates per sample), distributed on a 6,949 day-period (24 October 2001 to 2 November 2020) were analyzed (Fig. 1). The intervals between the samples ranged from seven days to 1,966 days/5.38 years (mean 331 days). Except for one sample from 2011 which yielded 10 isolates of a unique clone of *Hafnia alvei*, all other samples were positive for *E. coli stricto sensu* (49), leading to a collection of 210 *E. coli* isolates (Table S1).

## Phylogenomic analysis of the ED *E. coli* isolates and delineation of clones

### ED isolate core and pan genomes

The soft core genome (hereafter, “core”) of the 210 isolates (*i.e.*, genes present in 95% to 100% of the isolates) was composed of 3,278 genes. On the other hand, the pangenome (the totality of genes) encompassed 13,627 genes. The core and pangenome of the clones were similar to that of the isolates, although slightly smaller (3,270 and 12,986, respectively).

### Clonal diversity within the 210 ED E. coli isolates

*In silico* phylotyping, confirmed by a SNP based core genome phylogeny (Fig. S1), showed that all the known phylogroups (49), except the C phylogroup, were represented among the 210 isolates (Table 1). The most prevalent was the B2 phylogroup (n=72 isolates, 34.3%), followed by the A and F phylogroups (n=40, 19% each), the B1 phylogroup (n=26, 12.4%), the D phylogroup (n=17, 8%) and the H, G and E phylogroups (5, 1.9 and 0.5%, respectively). The most frequent STs were the STs 131 (20%, B2 phylogroup) and 59 (19.5%, F phylogroup) followed by the ST10 (11%, A phylogroup) (Table S2).

**Table 1.** Distribution of the 210 *E. coli* isolates gathered from the stools of the individual ED.

| Phylogroup | Nb (%) of isolates | Nb (%) of clones | Isolate/clone ratio |
| --- | --- | --- | --- |
| A | 40 (19) | 11 (31.4) | 3.6 |
| B1 | 26 (12.4) | 10 (28.5) | 2.6 |
| B2 | 72 (34.3) | 4 (11.4) | 18 |
| D | 17 (8) | 6 (17.1) | 2.8 |
| E | 1 (0.5) | 1 (2.8) | / |
| F | 40 (19) | 1 (2.8) | 40 |
| G | 4 (1.9) | 1 (2.8) | / |
| H | 10 (5) | 1 (2.8) | / |

For the delineation of clones, thirty four distinct haplogroups based on a combination of known typing markers (see methods for precise definition) were first identified (41). Then, the pairwise core genome SNP distance distribution was analyzed and confronted to the haplogroup of the isolates. It clearly appeared a cutoff separating inter-haplogroup isolates differing by more than 1840 SNPs from intra-haplogroup isolates differing by 21 SNPs or less (Table S3, Fig. S2). The only exception was in the phylogroup A_ ST399/661_O25:H12_fimHND haplogroup, where isolates differed by 125 to 130 SNPs. We checked the integrity of the mismatch repair genes in these isolates to exclude mutators (50). Based on our data, together with a recent modeling study defining genomic epidemiology thresholds from bacterial outbreaks between one to 19 SNPs (51), we considered that this haplogroup encompassed two different clones, named A and B. Nevertheless, we cannot exclude that all the isolates of this haplogroup belong to the same clone that evolved in the ED gut. In sum, our data-driven approach identified a maximum of 35 clones named thereafter “ED clones” among the 210 *E. coli* isolates (Table S4, Table 1). The main STs among the 35 clones were ST10 (11.4%) followed by ST131 and 69 (D phylogroup) (8.6% each).

The distribution of these clones in the phylogroups was very uneven. Excluding the rarely identified E, G and H phylogroup isolates, two categories of clones can be distinguished. On one hand, numerous clones are identified among the isolates (isolate/clone ratio from 2.8 to 3.6) in phylogroups B1, D and A, indicating a high genetic diversity whereas very few clones (isolate/clone ratio from 18 to 40) were identified in phylogroups B2 and F, respectively (Table 1).

### Global genomic characteristics and virulence, antibiotic resistance and antiviral defense island gene content of the isolates overtime

We first asked whether the global genomic characteristics of the isolates were constant in time. The mean of the total number of genes per isolate at a given day and the mean of the percentage of plasmid genome length among the complete genome were stable over years with a global mean at 4,768.27 and 2.30 %, respectively. The number of replicons per isolates was also relatively stable (global mean 3.83), with a clear predominance of Col and IncF plasmids (Fig. S3).

We then focused on specific genetic elements known to be important for *E. coli* commensal lifestyle in the gut: the virulence associated genes, as it has been proposed that virulence is a by-product of commensalism (52), the antibiotic resistance genes, as antibiotics are major selective agents and the defense islands against phage predation (53). These factors may be primarily colonization factors (4). We also compared the ED clones to an epidemiologically relevant control collection of 359 *E. coli* isolates from the stools of 359 healthy individuals living in France during the same period (54), to potentially identify ED-specific features.

When considering functional classes of virulence factors, i.e. adhesins, iron acquisition systems, protectins, toxins and invasins, we did not see any significant difference over time (Fig. 2A). Using random samples of 35 strains from the control commensal collection, we found the ED clones had significantly less virulence associated genes than the commensals (mean 47 in ED against 56 in commensals, p = 0.00014 according to a permutation test) (Fig. S4).

**Fig. 2.**
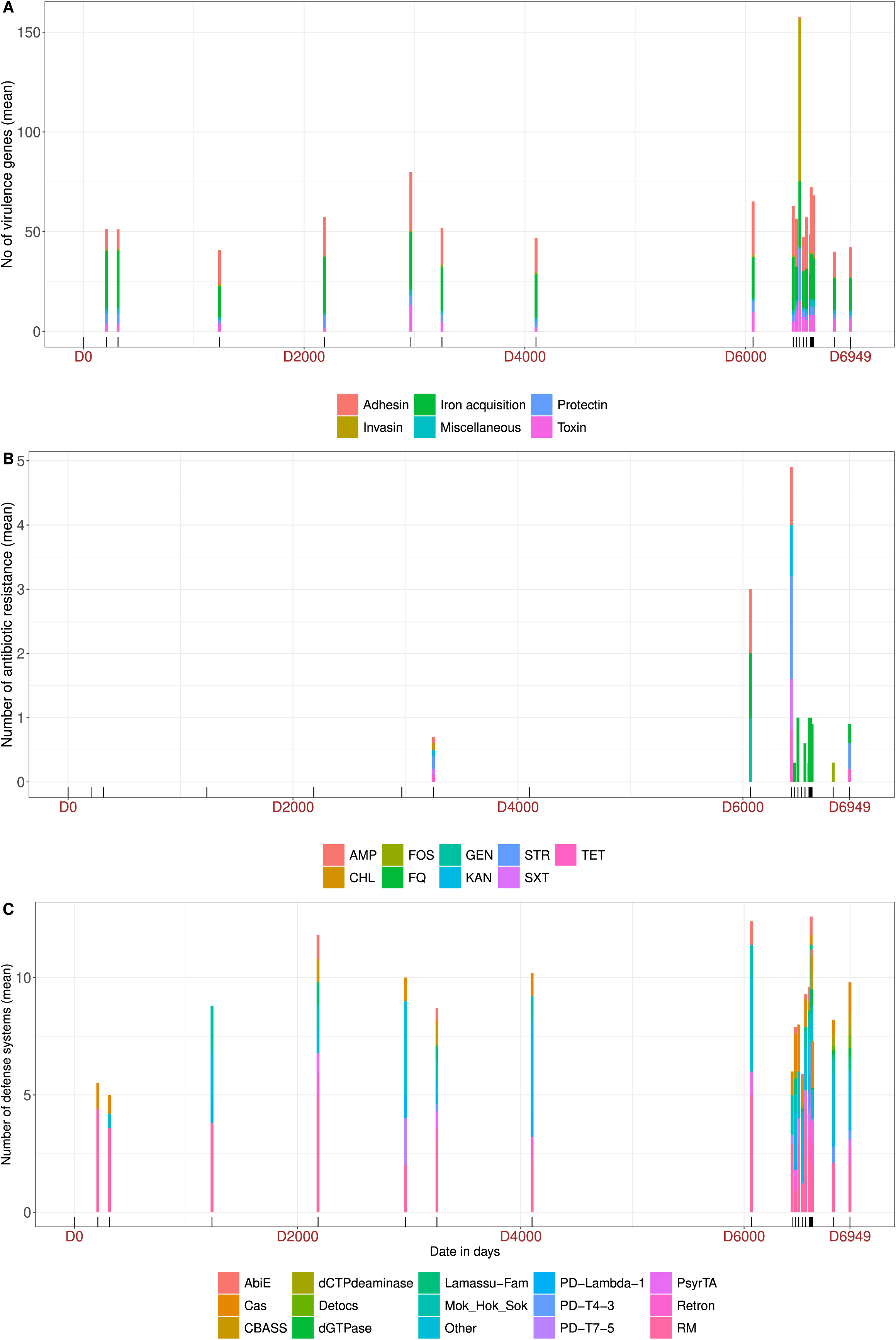
Mean of the cumulative number of (A) virulence associated genes, (B) antibiotic resistance and (C) antiviral systems per sample of the 210 *E. coli* isolates over time. At each date, the numbers of virulence associated genes, antibiotic resistance and antiviral systems of each of the 10 isolates are summed and the mean per isolate presented. (A) The genes are colored by main functional categories. Day 0 (D0) to day 6949 (D6949) are indicated in brown as in Fig. 1. (B) The phenotypic resistance pattern of the strains to main classes of antibiotics was deduced from the genotype as in (39). AMP: ampicillin, CHL: chloramphenicol, FOS: fosfomycin, FQ: fluoroquinolone, GEN: gentamycin, SXT: sulfamethoxazole trimethoprim, TET: tetracycline. (C) Antiviral systems colored by functional groups.

Genes or mutations conferring antibiotic resistance appeared briefly in 2010 in a ST362 strain exhibiting *bla_TEM1B_*, *sul2*, *tet(A)* and *catA* genes and were more prevalent starting in 2018, with five clones bearing resistance determinants: ST131 O16 (*bla_TEM1B_, aac*(*3*)*-IId*, GyrA 83), ST10 O89 (*bla_TEM1B_, sul1, sul2, dfrA5, tet(A)*), ST69 O23 (*bla_TEM1B_*), ST1086 (*fosA*), ST399 O129 (*tet(B)*), conferring resistance to ampicillin, aminoglycosides, quinolones, sulfonamides, fosfomycin, tetracyclines and chloramphenicol (Fig. 2B). No ESBL coding genes were detected. As the subject did not use antibiotics during the study period, these resistant strains were able to colonize in the absence of antibiotic selection. Using random samples of 35 strains from the control commensal collection, we found the number of resistance genes in the ED clones was slightly less than typical in commensals (mean 0.6 in ED against 1.1 in commensals, p = 0.045 according to a permutation test) (Fig. S4).

Finally, we searched exhaustively for the known defense systems in the genomes of the isolates using DefenseFinder (45). In all the 35 ED clones, we found 74 different defense system types (7.8 mean per bacterium +\-2.5 [3-13]). We did not see any trend over time for the number of defense systems, nor the diversity of defense (Fig. 2C). Using random samples of 35 strains from the control commensal collection, we did not find any significant difference in the diversity (number of different systems types) of defense systems with the ED clones (74 in total in ED against 69 in commensals, p = 0.87 according to a permutation test) (Fig. S4). However, two systems were significantly more abundant in the ED clones than in the control commensal strains (chi2 test corrected by Bonferroni): dGTPase and Detocs (Fig. S5). We will come back to this finding below in a more systematic analysis.

## Epidemiological dynamics of ED clones

We then investigated the epidemiological dynamics of the ED clones, which depend on how long clones persist in ED, and on the rate of colonization of the strains able to colonize ED.

### Residence times and dominance of ED clones

We quantified the residence time of ED clones with a Bayesian approach incorporating prior information on residence times measured in a previous study (48). We inferred the posterior distribution of the residence time for each observed clone (Table S4). Most of the clones (30) out of 35) were seen only once and consequently had very wide inferred posterior residence times distributions (with a mode at 2.1 days), reflecting the poor information on the true residence times of these clones in our data. The remaining five clones classified as resident have a posterior mode of residence time ranging from 10.1 to 437.1 days. Moreover, we classified clones by their dominance. Clones were considered dominant if present at least in one sample with at least 5 isolates; otherwise, they were classified as sub-dominant. Most of the clones (25 over 35) were sub-dominant with a mean of 2.1 isolates per sample. But most (15 over 21) timepoints have a dominant clone.

All in all, among the 35 clones, 25 were sub-dominant/transient (52 isolates), five were dominant/transient (48 isolates) and five were dominant/resident (110 isolates) (Table 2). Residence time was positively correlated with within-host frequency (linear model, effect size = 93 days, 95% CI [2.1-180], p = 0.045) (Fig. 3A). Phylogroups and STs were unevenly distributed among categories with B2 (STs 131 and 452) and F (ST59) phylogroups found in most of the dominant/resident strains but never as subdominant/transient. Conversely, phylogroup B1 (represented by numerous distinct STs), and to a lesser extent A and D, were found in subdominant/transient strains. Among the 21 samples, a dominant clone was found in 15 cases (71.4%) with in nine cases a full dominance (10 over 10 isolates) whereas in the six remaining samples, three to six clones were present per sample. The minimum duration of carriage of resident clones varied from seven days (ST635 O11) to 28 days (ST131 O25 B), 315 days (ST452), 489 days (ST59) and 1065 days (ST131 O25 C1), corresponding to inferred posterior modes of 10, 163, 234, 437 days respectively (Fig. 1).

**Table 2.**
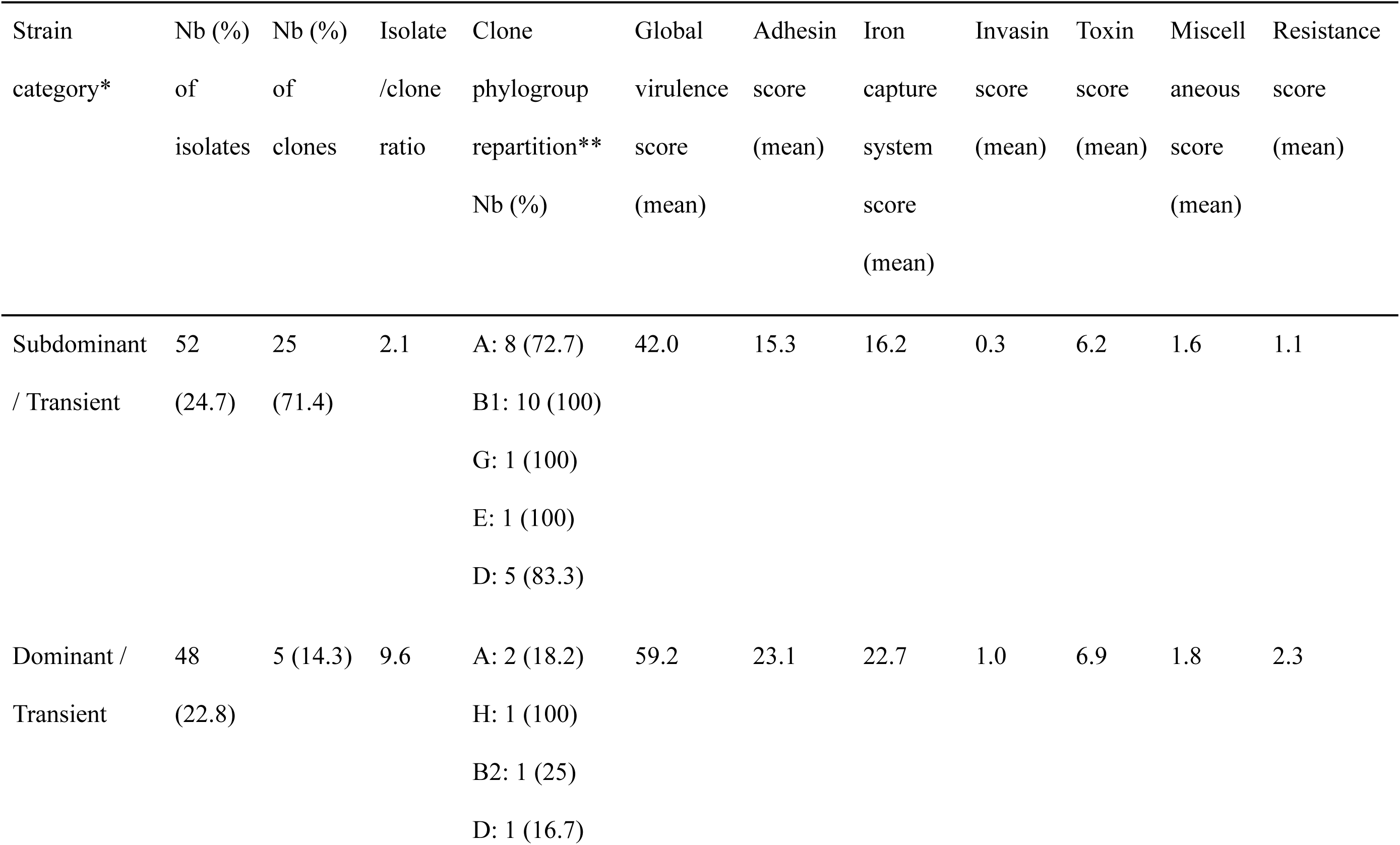
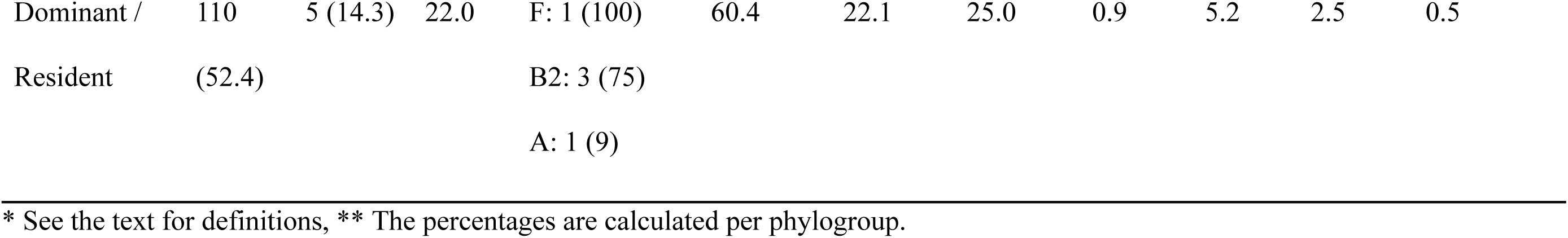
Feces carriage categories of the 210 isolates belonging to the 35 identified clones.

**Fig. 3.**
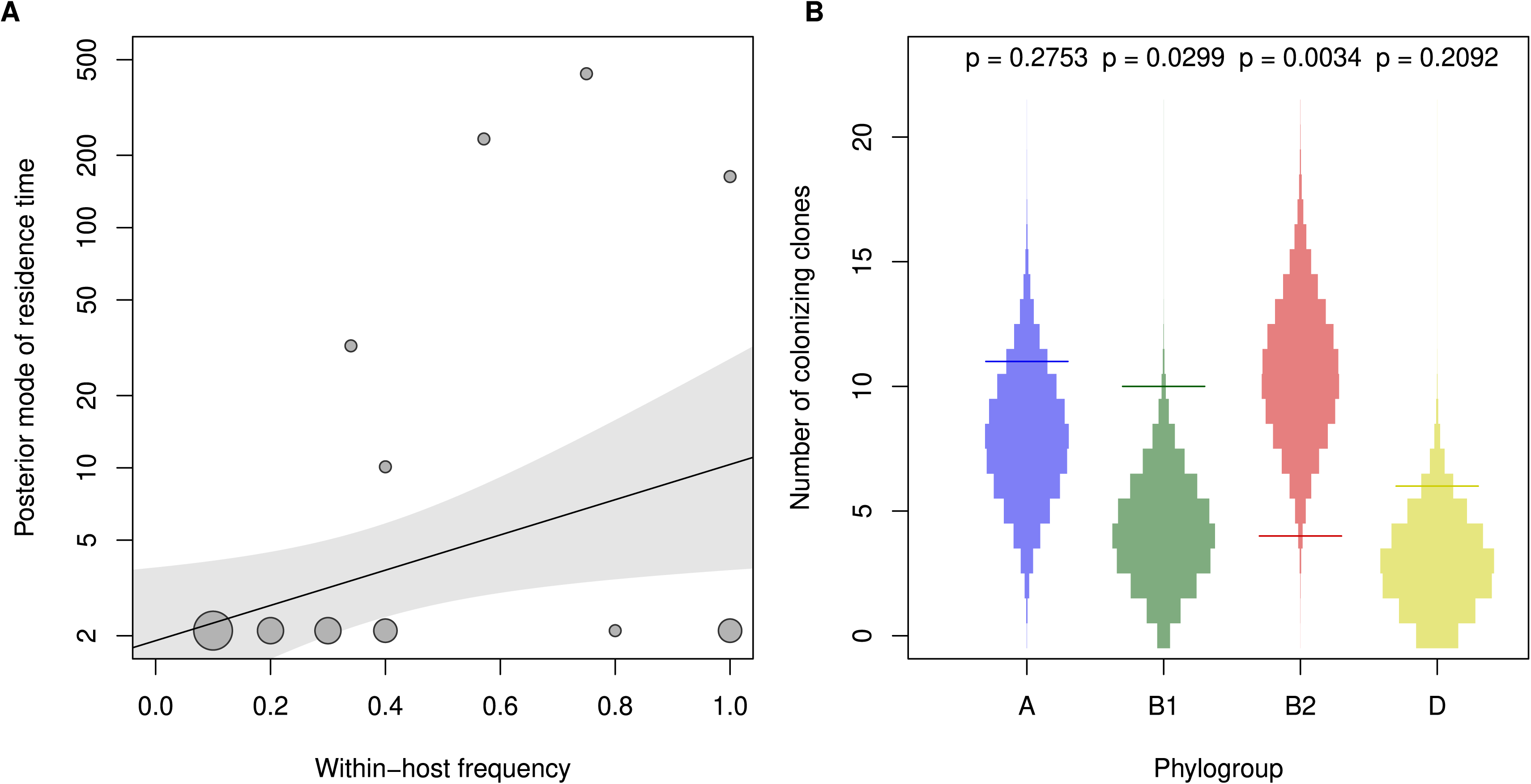
Residence time and colonization rates of the *E. coli* ED clones. (A) Inferred posterior of residence time of clones as a function of its within-host frequency (circles, area proportional to the number of observations), with the predicted relationship by linear regression (solid line, with gray area representing the 95% confidence interval). (B) Observed number of colonizing clones for phylogroup A (blue), B1 (green), B2 (red) and D (yellow) (horizontal line). This is compared to the expectation if ED was in contact with the representative pool of *E. coli* in the population (the 359 healthy individuals), (colored areas), used to generate p-values corresponding to the probability of these values under this hypothesis.

Long term intestinal residence of colonizing *E. coli* has been associated with the presence of specific clones belonging mainly to B2/D/F phylogroups, iron capture systems (17,55), and fluoroquinolone resistance (56). We did a series of tests for the association between the clone category, or inferred posterior mode of residence time, and various genetic factors. Although we found that clones with longer residence time exhibited more iron capture systems (Table 2), this association disappeared when controlling for the phylogroup.

### Colonization rates of ED clones and the trade-off with residence

The epidemiological dynamics of *E. coli* rely not only on the intestinal residence of clones, but also on their ability to colonize new hosts, which was assessed at the phylogroup level. The number of colonization events (i.e. the first appearance of a new *E. coli* clone) was computed for phylogroups represented by more than a single clone (A, B1, B2, D) and compared with the expected number of events assuming an equal colonization rate for all phylogroups, accounting for the relative phylogroup frequencies in France as estimated from the 359 *E. coli* control isolates from healthy individuals (54). Colonizations of the gut of ED by B1 and B2 phylogroups were respectively more frequent (4.72 more events than expected, p = 0.0301) and rarer (7.5 less events than expected, p = 0.0034), suggesting intrinsic differences in colonization rates (Fig. 3B). It therefore appears that the high residence time of B2 clones is counterbalanced by a lower colonization ability.

## Genome-wide genomic features associated with clones successfully colonizing ED

To identify distinct genetic features of *E. coli* strains able to colonize the focal individual, we performed a systematic genome-wide association study (GWAS) using Scoary (42) to investigate if some genes were differentially retrieved between the 35 ED pangenomes and the pangenomes from the 359 *E. coli* commensal strains of the control collection (54). The core genome of this control collection was composed of 3,262 genes whereas the pangenome encompassed 27,752 genes.

Indeed, we found 36 genes significantly associated with the 35 ED clones, of which 32 are enriched (rather than depleted) in ED clones (Table S5). Two clusters of biologically relevant genes were enriched in ED clones: a cluster of eight genes (*mhp*) allowing the use of the 3-(3-hydroxyphenyl)propionate (3HPP), one of the phenylpropanoids from lignin (57) and a cluster of 10 genes found in a phage defense island. In depth analysis of these two clusters is presented below.

### mhp cluster

This cluster is composed of the transcriptional activator gene (*mhpR*), enzyme coding genes *mhpA* to *F* and transport protein gene *mhpT* flanked by two bacterial interspersed mosaic element (BIME) (58). It was found in all 35 ED clones but only in 71.6% (257/359) of the control strains. An association with specific STs has been recently reported for this gene cluster (59), which is present in all phylogroups but not B2 where it is absent except in ST131. It is in fact also present in the B2 phylogroup ST452, the first ED clone, rendering all the ED B2 strains positive for this cluster and explaining its subsequent identification by GWAS.

To elucidate the phylogenetic scenario leading to such pattern, we searched for this cluster in 1,777 *E. coli* genomes of high quality from the RefSeq database, reconstructed the *mhp* phylogenetic history and compared it to the strain phylogeny. We first confirmed the presence of this cluster in all phylogroups except B2, where it is present in the ST131 and some single locus variants (ST131-like) (Fig. 4A and Table S6). We also found a *mhp* positive strain related to ST452 (ST3693, double locus variant), and six other *mhp* positive B2 strains from various and rare STs (ST646, ST7898, ST91, ST136, ST681, ST1155) scattered across the B2 phylogenetic tree (Fig. 4B). The entire *mhp* cluster phylogeny (Fig. 4A) is congruent with the strain phylogeny except for B2 strains (Fig. 4B). The *mhp* found in the latter can be divided into three main groups: the *mhp*-B2-a which gathers three strain pairs distributed throughout the tree; the *mhp*-B2-b found in highly divergent STs (ST646 and ST7898); the *mhp*-B2-c carried by ST131 strains (Fig. 4B). The physical map of the *mhp* cluster with flanking regions of representative *mhp* positive strains from phylogroup B2 (Fig. S6) showed an almost perfect synteny, i.e. conservation of genes order on the chromosome. Insertion sequences (IS*1,* IS*3* and IS*110*), are found in few genomes. But given their location, their diversity and the highly conserved position of the *mhp* gene cluster, we can probably rule out any role in the acquisition of this determinant.

**Fig. 4.**
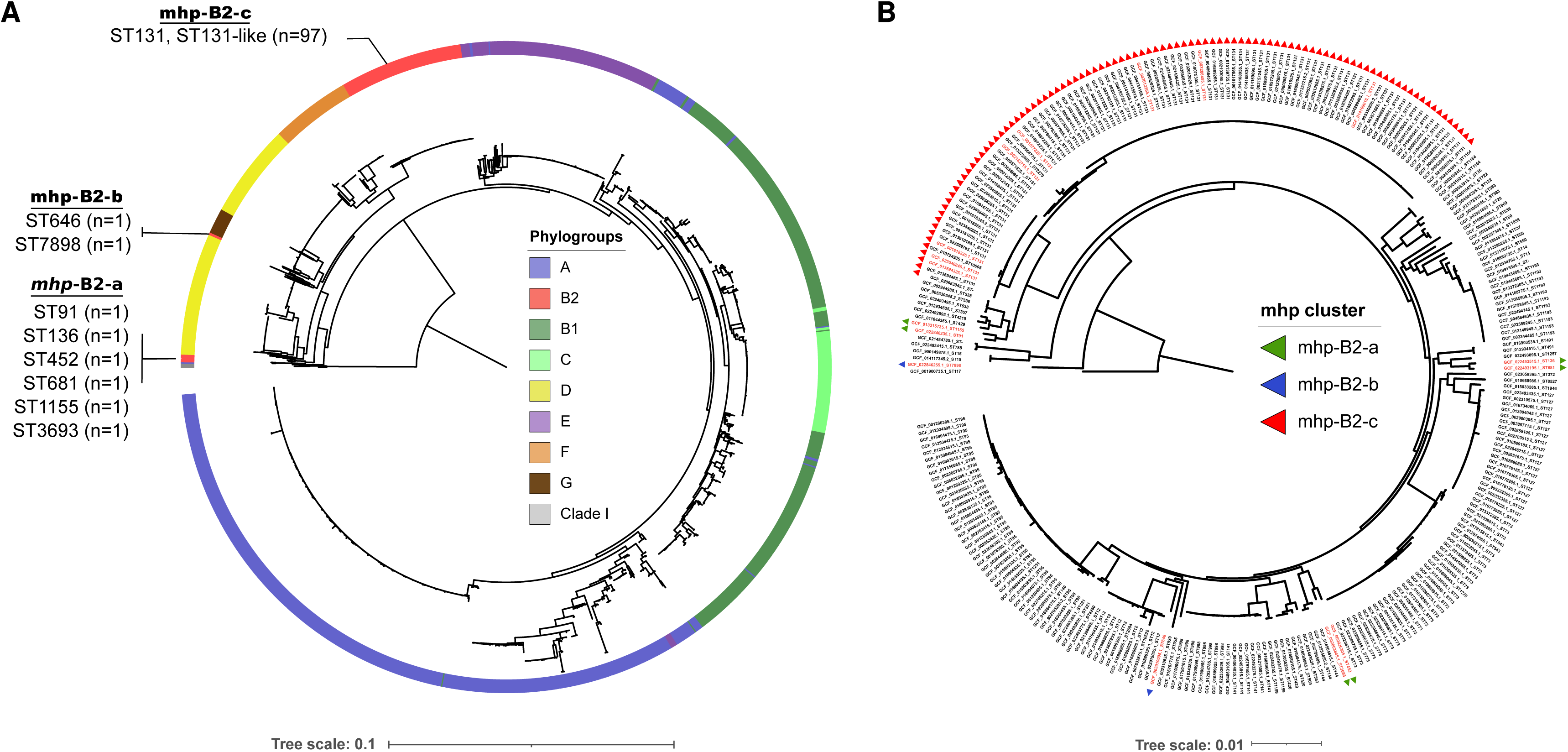
Phylogeny of the *mhp* gene cluster and its distribution among the phylogroup B2 phylogenetic history. (A) Maximum likelihood phylogenetic tree of the *mhp* gene cluster (7,884 pb) detected in 1576 over 1777 tested RefSeq genomes. The ED1a (NC_011745.1) *mph* gene cluster has also been included as no ST452 is available in the RefSeq dataset. The *mhp* carried by B2 strains is divided into three groups named *mph*-B2-a, *mph*-B2-b and *mph*-B2-c. The phylogroups are highlighted in color and the STs for the genomes belonging to phylogroup B2 are detailed. The tree is rooted on *Escherichia* clade I sequences. (B) Maximum likelihood phylogenetic tree computed from core gene alignment of the genomes belonging to phylogroup B2 in RefSeq genomes (n=282/1777). The genome of ED1a (NC_011745.1) has also been included. The presence of the *mph* gene cluster is highlighted by triangles colored according to the *mhp* groups previously defined: *mph*-B2-a (green), *mph*-B2-b (blue) and *mph*-B2-c (red). The tree is rooted on a phylogroup G ST117 genome (GCF_001900735.1). The genomes labeled in red correspond to the genomes used to analyse the genetic environment of *mhp* presented Fig. S6. Tree scales represent the number of nucleotide substitutions per site.

These data argue for an ancient arrival of the *mhp* cluster in the species followed by its loss in the ancestral B2 strain and several independent re-acquisitions probably by homologous recombination of flanking DNA regions as previously described for the High-Pathogenicity Island (HPI) (60). Another scenario, albeit less parsimonious, would be the presence of the cluster in the ancestral B2 strain, followed by multiple independent deletion events.

### Antiviral defense island

A cluster composed of 15 genes corresponding to an antiviral defense island (45), was present in five (5/35, 14.3%) ED clones (in complete and incomplete form in four and one clones, respectively) but never in the 359 control strains (Fig. 5A, Table S5 - genes highlighted in green). This island (18 kbp) was inserted between *pdeC-yjcB* and *uvrA-ssb* core-genes and thereafter named the *yjcB-ssb* defense island. It corresponds to the integration hotspot 39 (61). At least two defense systems were present inside the complete form: (i) a Detocs system composed of three genes, coding for a two-component system composed of an histidine kinase containing protein (DtcA), a “buffer” protein that absorbs phosphate signals (DtcB) and a response regulator (DtcC) (62). (ii) a *dGTPase* gene that encodes deoxyguanosine triphosphatase (dGTPase) enzyme, depleting the nucleotide pool and halting the phage replication (63). This gene is adjacent to (iii) *mazE*, coding for the antitoxin of the MazF/MazE module that suppresses phage lytic propagation via abortive infection (64). In the incomplete form of the island, Detocs genes are replaced by DNA mobility genes. In addition, one isolate/clone (ED2020-11-02-i) encodes for a different Detocs system (DtcC 48.1% similarity, DtcB 58.5% similarity, DtcA 37% similarity with Detocs system from ED2020-11-02-f), but in a different region of the genome.

**Fig. 5.**
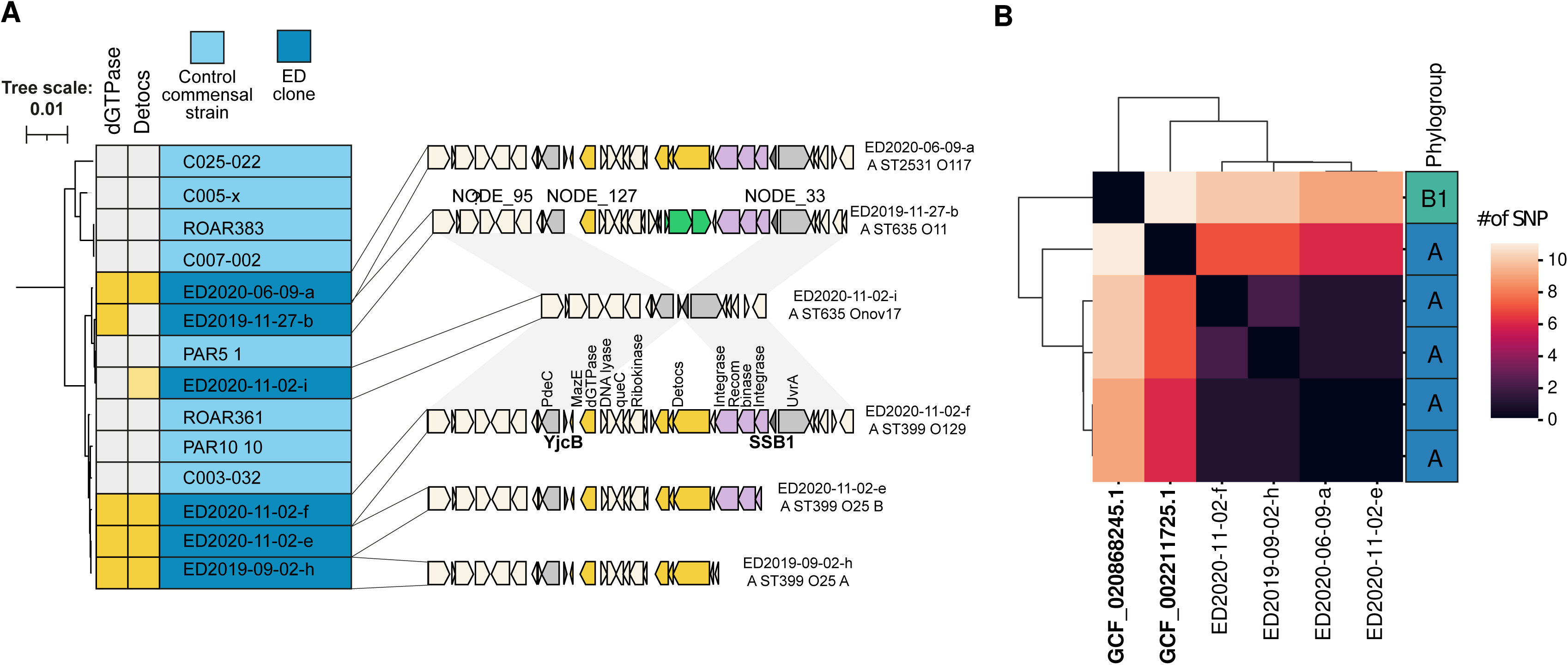
The *yjcB-ssb* antiviral defense island. (A) Schematic representation of the antiviral defense dGTPase and Detocs containing island found in complete and incomplete forms in four and one *E. coli* ED clones, respectively, with the synteny of the flanking regions in an isolate lacking this island (ED2020-11-02-i), but Detocs positive. Genes in yellow represent the different defense systems detected; in purple the mobility genes of the island; in green the insertion of another integrase inside the island; in gray the flanking genes *pdeC-yjcB* and *uvrA-ssb* genes. The island of the 2019-11-27-b isolate has been reconstructed using three different contigs (Nodes 95/127/33). The tree represents the phylogenetic history of the closely related phylogroup A0 strains belonging to the ED clone and control commensal collection based on the SNP core genome. Two main sub-lineages were observed, the upper branch corresponding to four island-negative strains whereas the lower branch corresponds to the Cplx399 with a mix of island-positive and negative strains. The presence or absence of dGTPase or Detocs is indicated in yellow. Of note, the ED2020-11-02-i isolate encodes for a different Detocs system (pale yellow), but in a different region of the genome and without the dGTPase. Conversely, the ED2019-11-27-b isolate is only dGTPase positive. (B). Clustermap based on the SNP matrix of the complete *yjcB-ssb* island (18 kbp) from the RefSeq database (in bold) and the ED clones. The A phylogroup strain from RefSeq belongs to the ST 635 whereas the B1 strain belongs to the ST58.

To characterize the phylogenetic context of the six clones carrying the defense island and/or Detocs system, we mapped these strains on a core genome tree reconstructed using the 35 ED clones and the control commensal strains. The *yjcB-ssb* defense island positive strains belonged to two closely related lineages (ST635/2531 and ST399), also called ST399 Cplx, within the phylogroup A0 (*arpA* +, *chuA* -, *yjaA* -, TspE4.C2 -) (65). They are intertwined with the ED2020-11-02-i isolate and with island-negative control strains (Fig. 5A). When searching for this island in the *E. coli* genomes from the RefSeq database, we found only two positive strains for the complete island belonging to phylogroup/ST A ST635 and B1 ST58 (Fig. 5B). SNP comparison of 12 kbp of the defense island (present in a single contig in all clones) between the different isolates showed very few SNPs among the ED clones (maximum 2 SNPs) (Fig. 5B), but a little more between the islands in the six ED clones and the island in the RefSeq A ST635 strain (between 7-8 SNPs), the B1 strain island being more phylogenetically distant (between 9-11 SNPs).

The *yjcB-ssb* defense island was observed for the first time in September 2019 (day 6522) and found subsequently in November and December 2019 (days 6608 and 6615), and June and finally November 2020 (days 6803 and 6949) (Fig. 1). If we consider individually the two main defense systems, the ST635 O11 and ST635 O17 clones are only dGTPase and Detocs positive, respectively.

## Within-host evolution of resident ED clones: no evidence for diversifying selection and estimation of the mutation rate

In previous work studying the three first samples of the time series, we analyzed the genomic diversity of the ST452 clone at the SNP level using Illumina technology coupled to a circularized genome of high quality (ED1a) (24,29). We pursued this analysis on nine additional clones (128 isolates) (Table S7) corresponding to unique samples in five cases where the clone was dominant (8 to 10 isolates per sample) and to four clones (8 to 40 isolates per clone) sampled at multiple dates, with seven to 1065 days separating the first and last samples. We used Illumina technology (short reads) on all isolates associated with one isolate sequenced with Nanopore technology (long reads) per clone. Using breseq 0.33.2 (47) we identified 48 different mutations, 43 of them SNP and 33 of them genic SNP. No parallelism was observed at the gene level, as all the mutations targeted distinct genes (Table S7). The ratio of non-synonymous-to-synonymous mutations was estimated, based on 20 codon usage frequency tables established for *E. coli* from international DNA sequence databases (66) including more than 1,000 codons. The average ratio was 0.836 (from 0.730 to 0.889), consistent with weak purifying selection.

The observed mutation rate of *E. coli* in the ED samples was then estimated as the slope of a regression of the number of new mutations accumulated by each clone over time (67), at 1.503×10^-9^ mutation per day per nucleotide (95%CI=[1.159×10^-9^, 1.848×10^-9^]), i.e. 5.491×10^-7^ mutation per year per nucleotide or 2.77 mutations per year per genome (Fig. 6A). This observed rate does not include the highly deleterious mutations which cannot be observed in our approach. Based on previous estimations of the mutation rate at 2.2×10^-10^ mutation per generation per nucleotide (67), the estimated generation time of *E. coli* in the ED samples is therefore 3.51 hours (95%CI=[2.86, 4.56]). Given this generation time, a single ingested *E. coli* could reach densities of the order of 10^8^ cells/g of feces in around five days (128.3 h), assuming exponential growth and a colon holding 1 kg of feces. Assuming neutrality, only the mutation rate and the effective size (*N_e_*) of the *E. coli* population should define the divergence between two isolates of the same clone over time. Using a stochastic model describing the appearance and fixation of mutations, we simulated the number of mutations separating two initially identical genotypes over time, for the estimated mutation rate and various effective population sizes. The divergence observed in ED was consistent with a value of *N_e_*of the order of 1000 bacteria, the model predicting an equilibrium reached in about a year (Fig 6B). For lower effective sizes (e.g. *N_e_* = 100), the divergence stabilized at a much lower equilibrium than observed, while for larger ones (e.g. *N_e_* = 10000), it continued increasing after a year towards much higher values.

**Fig. 6.**
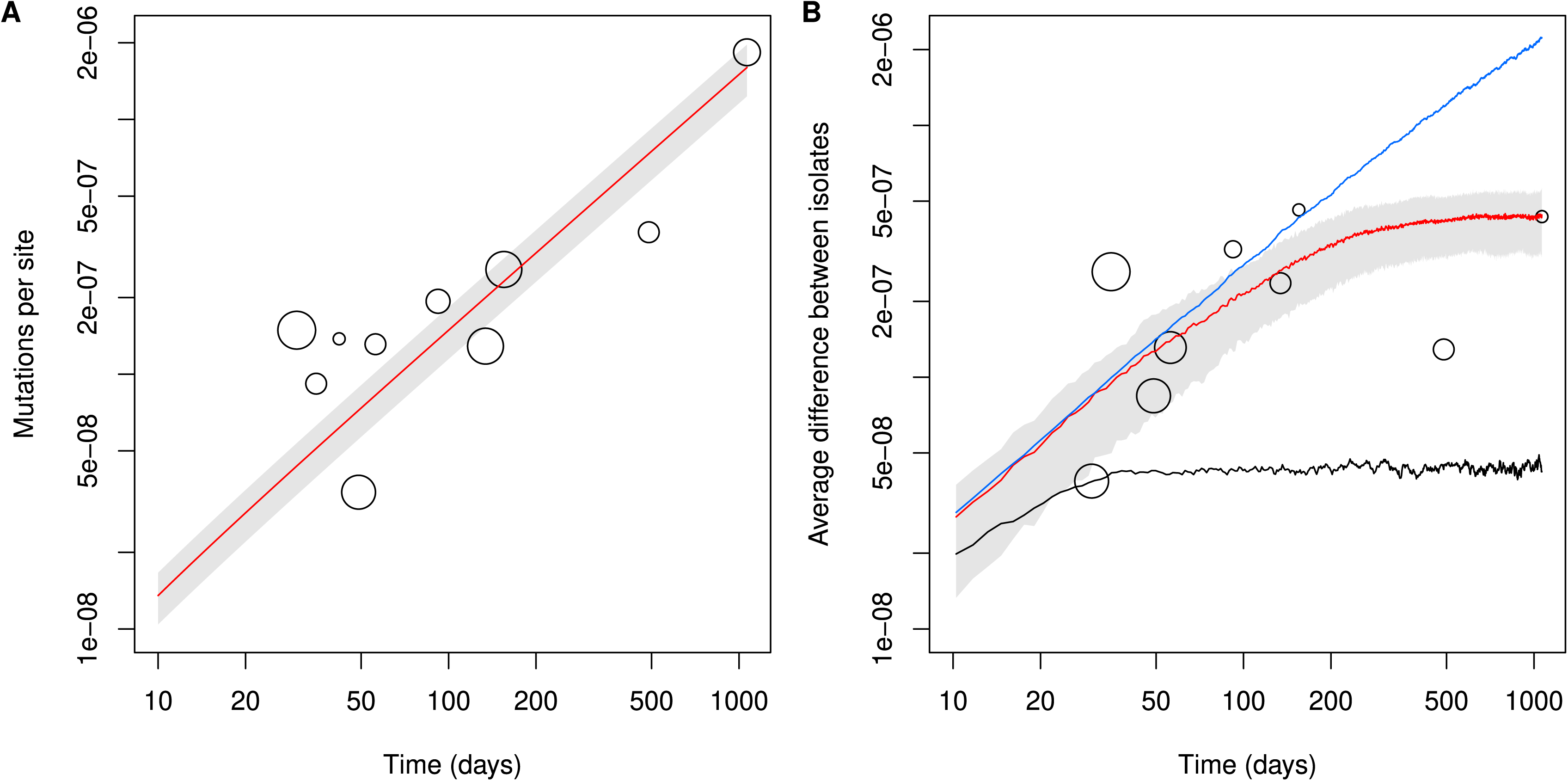
Accumulation of mutations over time in the *E. coli* ED clones. (A) Number of observed mutations per site (in log-scale) as a function of time elapsed since the first sampling (in log-scale), for each sampling of the four clones sampled multiple times after the first one (80 isolates, circles, size proportional to the number of isolates), with the estimated observed mutation rate (red line) with the predicted 95% confidence interval (gray). (B) Average difference between isolates of the same clone per site (in log-scale), as a function of time since the first sampling (in log-scale), for each sampling of the nine clones considered (circles, size proportional to the number of isolates) (Table S7), and averaged over 1,000 simulations of the Wright-Fisher model with *N_e_* = 100 (black line), with *N_e_* = 1000 (red line with interquartile range representing 50% of the simulations in gray) and with *N_e_* = 10000 (blue line).

## Discussion

This work is based on the originality of our collections with (i) the strains from the ED individual that can be considered as “wild” commensals of a high income country due to the absence of medical records of the host, as confirmed by the low level of virulence and resistance genes and (ii) the commensal strains from the French subjects that were used as a control cohort. The most striking features are discussed below.

### Large diversity of *E. coli* hosted by a single individual

Within a single individual, we observed a large diversity of STs, and all phylogroups were represented. As a consequence, the soft core genome of *E. coli* in a single individual (n=3,270), was almost identical to the control commensal collection of trains from 359 individuals (n=3,262). When additionally comparing the ED core genome to a more diverse datasets chosen to reflect the diversity of the whole species, represented by 1,294 Australian strains from various origins (mammal and bird hosts, environment) selected from more than 5,000 strains, we also found near identical core genomes (n=3,242) (68) (data not shown). The pangenome was composed of more than 13,000 genes. Considering that the human genome encompasses 20,000 protein-coding genes (69), this means that the individual hosted a number of *E. coli* genes representing almost 70% of the number of his own genes.

### The gut colonization/residence phenotypes are phylogroup/lineage dependent

We were able to demonstrate two paces of strain turnover in the human gut: rapid turnover with a timescale of the order of one week, for highly diverse B1 and most A clones (sub-dominant and transient phenotypes); and slow turnover with a timescale of the order of one year, for a few extra-intestinal pathogenic *E. coli* belonging to B2 (ST131) and F (ST59) phylogroups (dominant and resident phenotypes). The longest observed period of carriage was three years (ST131 O25 C1). We pursued the ED stool analyses in an ongoing research project and found the ST59 strain as dominant in the stools of ED in December 2021, corresponding to 2 years and 5 months (884 days) of asymptomatic carriage (70).

The pattern of coexistence of phylogroups within the single ED subject was in agreement with what was previously found in a large cohort of 100 healthy French subjects with 100x higher threshold of detection of the four main phylogroups (71). The authors showed that, when phylogroup B2 was dominant, it tended not to co-occur with other phylogroups. In contrast, other phylogroups were present when phylogroup A was dominant. The dominant B2 clones belonged mainly to archetypal ExPEC ST95, 131 and 73 (49).

The link between B2/D phylogroups and long residence time in the gut has been reported in analyses of a longitudinal study of 70 Swedish infants based on the identification of the four main phylogroups (72). Using the more complete scheme of the Clermont typing method, Martinson *et al*. showed in a smaller and more detailed longitudinal cohort study (8 adult participants, 8 to 30 months) that A (corresponding mainly to ST10 complex), B2.3 (corresponding to the majority of B2 ExPEC) and F phylogroups are more likely to be human residents compared to the other genotypes (48). Similarly, longitudinal cohort study of neonatal gut microbiomes showed that ExPEC clones (ST73, 131, 69, 95 and 59) were the best gut colonizers within the first 21 days, tended to dominate the *E. coli* flora and could reside up to one year (73). A re-analysis of the data from Martinson and Walk (48) and a study on a cohort of 46 healthy French adults (70) showed that the long residence of some phylogroups was counterbalanced by a poorer capacity to colonize hosts. This residence-colonization trade-off, together with an additional mechanism of niche differentiation, could in theory explain the coexistence of multiple *E. coli* phylogroups in the population (70).

As there is a link between B2 strains and the abundance of virulence genes (74), it is difficult to untangle the respective roles of the phylogenetic background and virulence genes. A recent study on 222 American veterans and household members (17) showed that the ST131-O25-*fimH30* was the strongest univariate correlate of intestinal residence. However, in the multivariate analysis, multiple virulence genes (especially the iron uptake systems) and fluoroquinolones were better predictors of residence times than the ST131 background. Our data, possibly because of the small sample size, were also non conclusive. Larger datasets on the duration of residence in the gut, together with whole genome sequences, would enable a GWAS that may be powered to untangle effects of individual variants from the genetic background.

A key unanswered question is how such diverse B1 and at a lesser extent A and D phylogroup strains can colonize the ED stools (often two different clones per sample) knowing that carriage of B1 strains in humans in France is rare (13%) (75), B1 strains being commonly found in cattle (76) and also in water and sediments (77).

## Host specific traits in the *E. coli* genomes: role of metabolism and phage predation

Beside the genomic diversity of the ED clones, we identified genes significantly enriched in the individual as compared to a pool of 359 individuals living in similar conditions. These genes are involved in two fundamental processes of gut colonization: metabolism (78) and anti phage defense (53). They are localized on mobile genetic elements and selected together with the clones bearing them.

The gene cluster encoding the 3-(3-hydroxyphenyl)propionate (3-HPP) catabolic pathway is enriched in ED strains, particularly in resident B2 strains from ST452 and ST131. This pathway allows strains to use aromatic acids such as phenylpropanoids as sole sources of carbon for growth. Such compounds are the result of the action of intestinal microflora on cereal constituents (79). It is tempting to speculate that the enrichment in *mhp* genes in ED is linked to his high and daily consumption of cereals (see SOM). Further microbiota studies should be performed in relation to diet to test the hypothesis. The *mhp* cluster, invading the *E. coli* species probably by horizontal gene transfer (58), is adjacent to the lactose operon (Fig. S6), both sets of genes allowing catabolism of breakfast compounds, i.e. milk and cereals.

Bacteria protect themselves from foreign DNA as phage DNA by more than a hundred systems, unveiling many different molecular mechanisms (53). We showed here that six clones belonging to the ST399 Cplx and bearing the very rarely reported dGTPase and/or Detocs defense systems are present during the last year of the study, suggesting that these clones are selected by a phage sensitive to the Detocs and/or dGTPase system(s) within the fecal phageome of ED individual. Indeed, although closely related, these clones came from distinct colonization events as attested by the number of SNPs between them (140 to 10,000, Table S3). However, no evidence on which phage(s) may be involved is available in the literature (80). Moreover, the rarity of this island suggests strong selection at the between-host level. The island is present in none of the 359 control strains, and in two out of 1,777 high-quality *E. coli* genomes. The frequency of the island in the natural *E. coli* population could thus be of the order of 0.1%. The occurrence of six events of colonization with strains carrying this island implies that at least ∼6000 *E. coli* strains contended to colonize ED over the 14 months when the six strains successfully colonized (Sep 2019 - Nov 2020), resulting in a rate of at least ∼14 *E. coli* strains per day contending to colonize a host.

These observations suggest that “Everything is everywhere, but the environment selects” (81). Individual hosts constantly ingest large numbers of diverse contending *E. coli* strains. Only a small fraction of them successfully colonize. Successful colonization can result from the adaptation of specific genotypes to the particular environmental conditions in the gut at this time.

## No major sign of selection over time within the gut of a single individual

It has been reported that *E. coli* isolates are adapting under strong selective pressure when colonizing an extra-intestinal site. This was based on genomic signs of selection, such as a high nonsynonymous to synonymous mutation ratio and mutational convergence (82). At the opposite, no molecular footprint of selection has been identified in commensal *E. coli* isolates from healthy individuals (83), including the previously published ST452 clone isolated from ED (24). In this study, we did not evidence a strong pattern of diversifying selection as in extra-intestinal sites, but weak purifying counter-selection of deleterious mutations. Nevertheless, we note that one clone splitted into two haplotypes separated by six mutations (see SOM), possibly by diversifying selection, but the wider significance of this single observed event was unclear. We acknowledge though that clear signatures of within-host selection could possibly be detected in larger datasets identifying more resident strains. Besides, we confirmed that levels of diversity are compatible with an effective population size much lower than census size, possibly due to population structure within the gut.

## Conclusions

Despite originating from a single individual, our data strongly suggest that the normal *E. coli* microbiota is shaped by both the intrinsic properties of clonal lineages and environmental constraints such as nutrient availability and presence of phages. Although we evidenced selection at the between-host level, no major sign of within-host molecular adaptation was identified. Such long term time series studies of *E. coli* microbiomes in a large range of subjects worldwide should be supported to generalize our findings.

## Supporting information

MaMeS

## Acknowledgments

We are grateful to Julie Marin for providing the control commensal data sets. We thank Guillaume Achaz for pointing out the method to analyze SNP diversity in longitudinal samples.

## Data Availability Statement

The datasets generated during and/or analysed during the current study are available in ENA database on the bioproject PRJEB45090 (https://www.ebi.ac.uk/ena/browser/view/PRJEB45090).

## Supplementary online material (SOM)

### Supplementary Tables

Table S1. Main characteristics of the 210 *E. coli* ED isolates

Table S2. ST distribution of the 210 *E. coli* ED isolates

Table S3. SNPs matrix of 210 *E. coli* ED isolates

Table S4. Main characteristics of the 35 *E. coli* ED clones

Table S5. List of genes significantly associated to the 35 ED clones

Table S6. Distribution of *mhp* gene cluster among 1,777 complete *E. coli* genomes from RefSeq

Table S7. List of mutations identified in nine *E. coli* ED clones corresponding to 128 isolates

## Supplementary Figures

**Fig. S1. Maximum likelihood phylogenetic tree reconstructed from the core genome single nucleotide polymorphisms (SNPs) (n=3 217 191) of the 210 *E. coli* ED isolates.** The three isolates of the ST399 O25B (ED2020-11-02-e and ED2020-11-02-h) and ST399 O25A (ED2019-09-02-h) are indicated by red and blue arrows, respectively.

**Fig. S2. Distribution of the pairwise core genome SNP distances according to whether pairs of isolates are inter- or intra-haplogroup.**

For clarity, the SNP scale is logarithmic with 0.1 for 0. The red dotted line corresponds to the cutoff chosen for the definition of clones. The dots circled in red correspond to the phylogroup A_ ST399/661_O25:H12_fimHND haplogroup isolates differing by 125-130 SNPs.

**Fig. S3.** Mean of the cumulative number of replicons per sample of the 210 *E. coli* ED isolates over time. A, B and C panels correspond to the year, month and week scale of sampling, respectively, as in Fig. 1.

**Fig. S4. Distribution of resistance, virulence genes and anti-phage defense systems between ED clones and commensal strains.** Mean number of resistant and virulence genes, and total number of defense genes in the 35 ED clones (vertical lines), together with the null distribution when sampling 35 strains at random among the 359 commensals of the control collection (histograms). The ED clones count slightly less resistant genes and less virulence genes, but a similar number of defense systems. The sampling of the control collection was done 50,000 times.

**Fig. S5. Percentage of strains encoding each defense system between the ED clones and the control commensal collection.** Only two systems are significantly more abundant in the ED clones (chi2 test corrected by Bonferroni, significance threshold p < 0.05): dGTPase and Detocs.

**Fig. S6. Physical map of the *mhp* cluster and its genetic environment among phylogroup B2 representative genomes.** Representative genomes from phylogroup B2 were selected from the RefSeq dataset for each *mhp* environment based on nucleotide clustering (identity=99%, coverage=95%). The ED1a ST452 genome (GCF_000026305.1) was also included. White arrows represent the *mhp* gene cluster. Genes that are homologous between different genomes are the same colour. The links between the maps represent the identity of the proteins. Insertion sequences are highlighted in red (IS*1*), blue (IS*3*) and green (IS*110*). The combination of serotypes and *fimH* allele was used to assign genomes belonging to ST131 to the clades A (O16:H5*/fimH41*, B (O25B:H4/*fimH161,* O25B:H4/*fimH22*) or C (O25B:H4/*fimH41*).

