## Supplementary figures and images for "Strain intrinsic properties and environmental constraints together shape *Escherichia coli* dynamics and diversity over a twenty-year human gut time series"

### FigS1

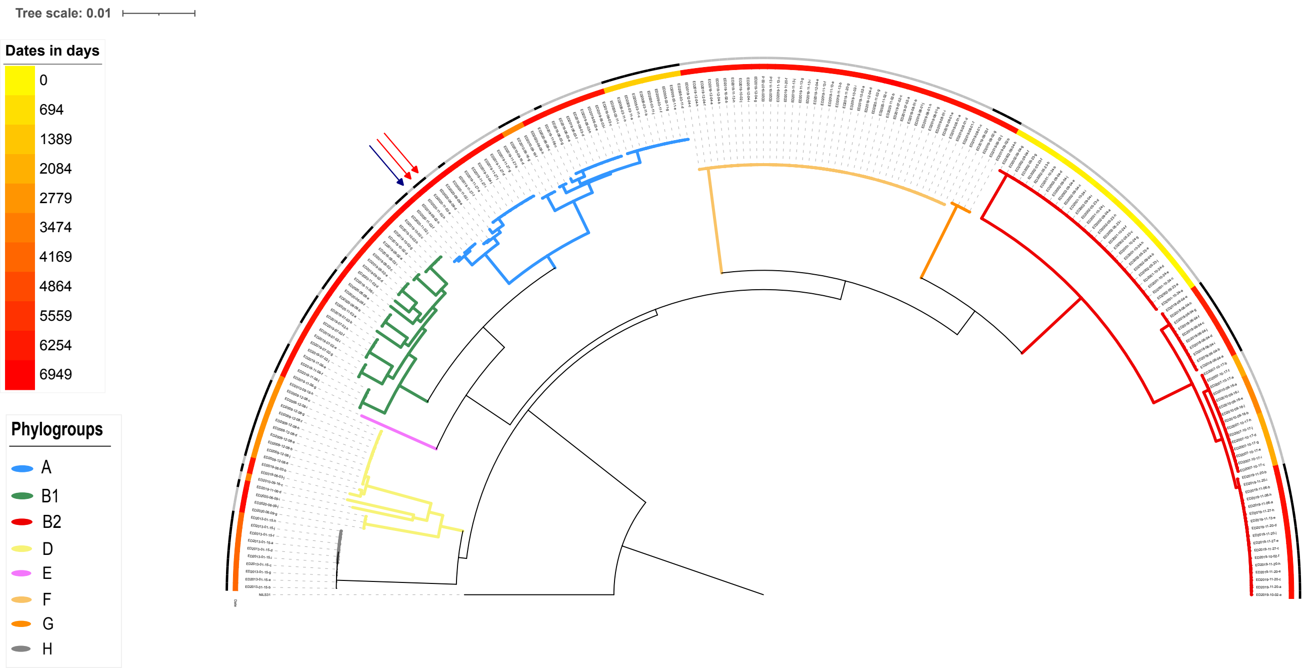

### FigS2

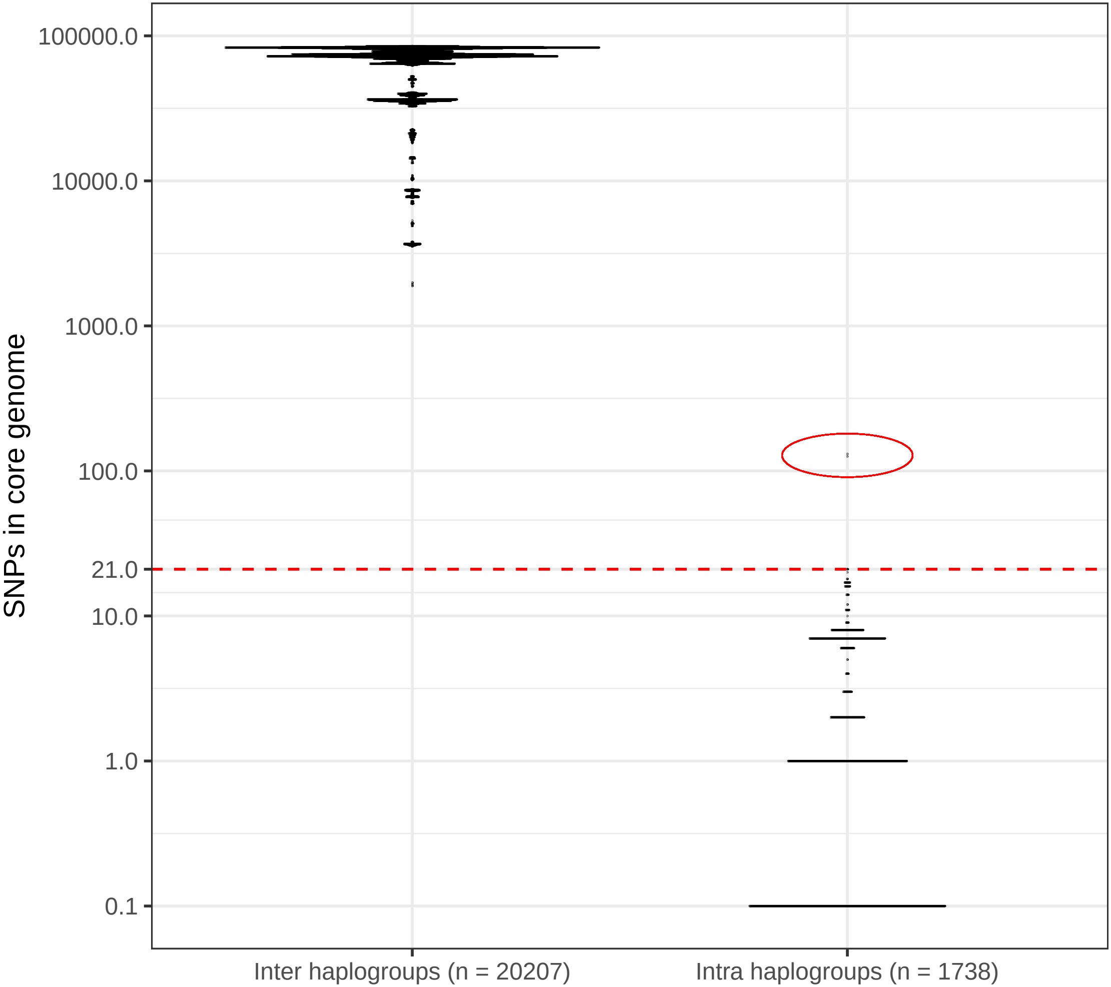

### FigS3

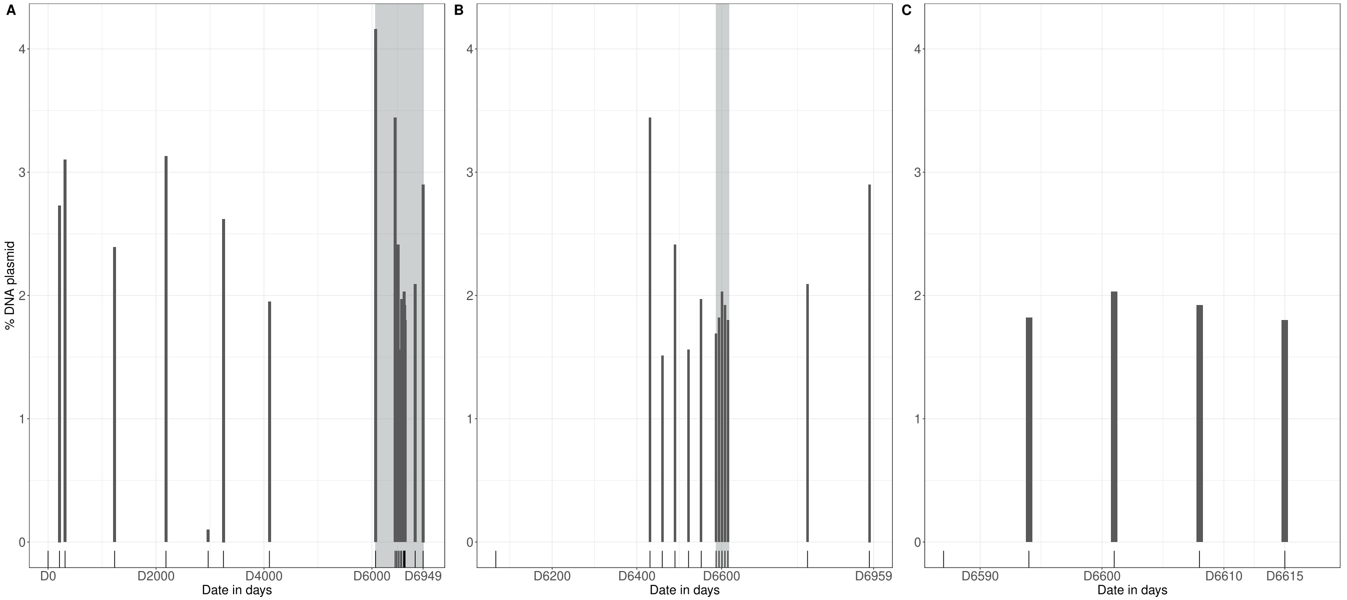

### FigS4

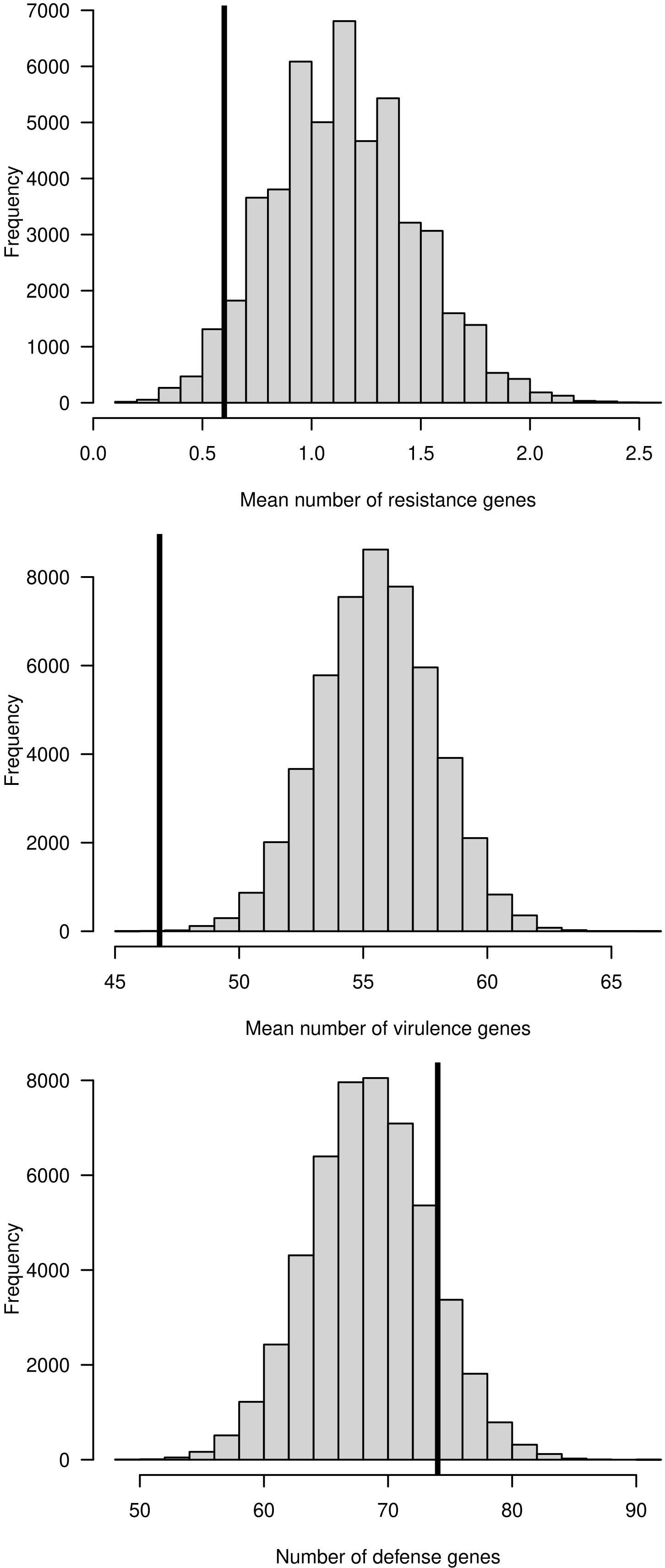

### FigS5

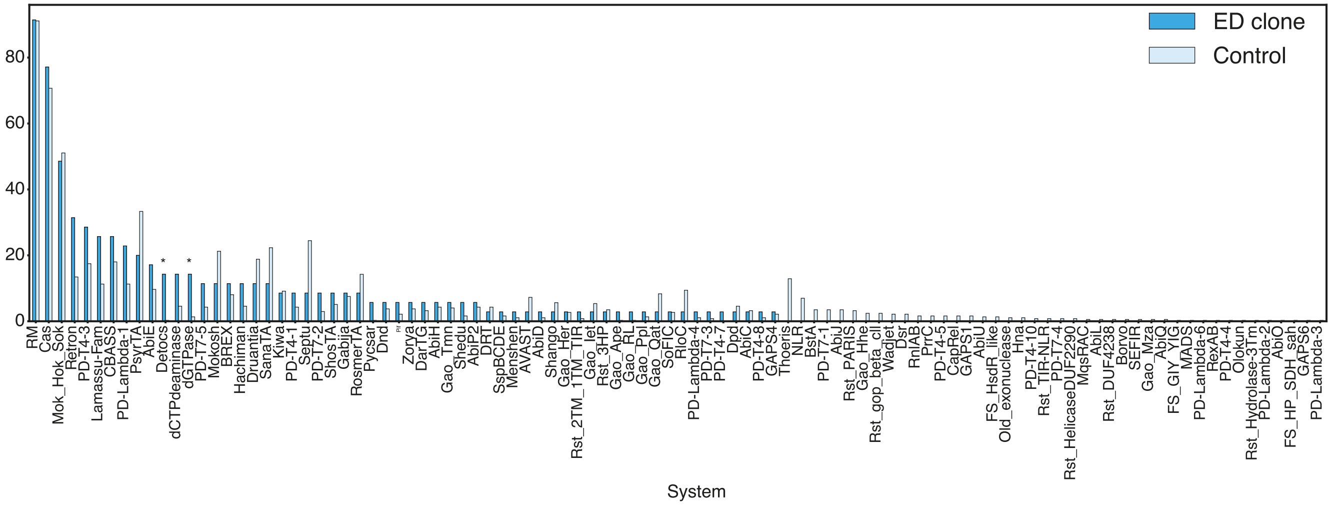

### FigS6

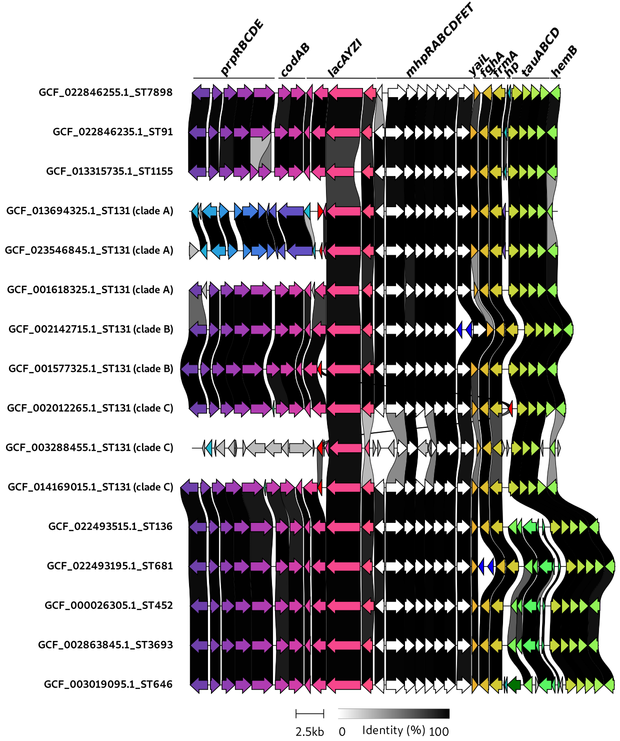
