## Supplementary material for "Strain intrinsic properties and environmental constraints together shape *Escherichia coli* dynamics and diversity over a twenty-year human gut time series": MaMeS

**The following presents supplementary online material (SOM) of the article "Strain intrinsic properties and environmental constraints together shape *Escherichia coli* dynamics and diversity over a twenty-year human gut time series" by:**

Bénédicte Condamine[1], Thibaut Morel-Journel[1,2], Florian Tesson[1, 3], Guilhem Royer[1,4,5], Mélanie Magnan[6], Aude Bernheim[3], Erick Denamur[1,7*], François Blanquart[8*], Olivier Clermont[1*]

[1] Université Paris Cité, INSERM, IAME, 75018 Paris, France

[2] Université Sorbonne Paris Nord, INSERM, IAME, 93000 Bobigny, France

[3] Institut Pasteur, Université Paris Cité, INSERM, Molecular Diversity of Microbes Lab, 75015 Paris, France.

[4] Unité de Bactériologie, Département de Prévention, Diagnostic et Traitement des Infections, AP-HP, Hôpital Henri Mondor, Créteil, France.

[5] EA 7380 Dynamyc, EnvA, UPEC, University of Paris-Est, Créteil, France

[6] Université Paris Cité, INSERM, CNRS, Institut Cochin, UMR 1016, 75014, Paris, France.

[7] AP-HP, Laboratoire de Génétique Moléculaire, Hôpital Bichat, 75018 Paris, France

[8] Center for Interdisciplinary Research in Biology, CNRS, Collège de France, PSL Research University, 75005 Paris, France

* ED, FB and OC shared last authorship on this work

### Material/Subject and methods

*Description of the ED subject*

ED stands for the initials of the subject. ED is a caucasian man, married with children, living and working in Paris. He was 44 year-old at the beginning of the study, weighing around 65 kg for a height of 1.75 m. Importantly, he did not take any antibiotics nor had been hospitalized during the sampling period. He is omnivorous with few (twice a week) meat and fish consumption, regular (daily) vegetable and fruit consumption and enrichment in dairy products and cereal (every breakfast) intake. He drinks a cup of wine every day. In a simplified dietary questionnaire designed to characterize the diet quality of the French population [1], ED scored the highest for cereal consumption (1/day) and dairy products (1/day). He traveled on average once a year outside France (Europe, USA, China, Australia) for short periods (less than two weeks). He never had a pet.

In sum, ED can be globally considered as representative of the healthy French urban adult population, with some diet specificities.

*Stool plating and isolate sampling*

Fecal samples were stored at 4°C after emission and transmitted to the laboratory. An aliquot of the fresh feces (one inoculation loop) was spread on Drigalski agar plates pure or sterile water diluted at $10^{-2}$ to obtain at least 10 isolated colonies. The plates were incubated at 37°C overnight (O/N), and the feces were discarded. Ten colonies (called “isolate” in the main text) with a yellow appearance were isolated from each plate at each time point. These colonies were then grown on liquid lysogeny broth (LB) O/N and stored at −80°C in glycerol. We choose to study 10 colonies because this level of depth is often used in the literature and it represents a good trade-off between the detection of diversity and the amount of work and money spent.

yjcB-ssb *defense island and* mhp *gene cluster detection and analysis*

The *yjcB-ssb* defense island detection within 1,777 RefSeq complete genomes, was based on the detection of both flanking genes of more than 300 bp of the island (*pdeC* and *uvrA* genes).

We searched for the presence of the *mph* gene cluster in the 1777 RefSeq genomes using a custom database for abricate (https://github.com/tseemann/abricate) with identity and coverage thresholds of 90%. We considered the presence of the cluster only in genomes carrying all genes (i.e. 8 genes : *mhpRABCDEFT*). Then, sequences were extracted using samtools v1.18 [2] and aligned with mafft v7.490 [3]. We computed a maximum likelihood phylogenetic tree from the alignment using iq-tree v1.6.12 [4] with the option "MF" to select for the best-fit model (i.e. TIM3+F+R2).

We then compared the evolutionary history of *mhp* with that of the B2 phylogroup strains. In brief, we constructed a pangenome using Ppanggolin [5] including all RefSeq B2 strains (n=282/1777), ED1a (NC_011745.1) and a genome belonging to phylogroup G (GCF_001900735.1) as outgroup. Then, we computed a maximum likelihood phylogenetic tree from core gene alignment of these 284 genomes with iq-tree with the option "MF" to select for the best-fit model (*i.e.* GTR+F+R4). We determined multi-locus sequence types using MLST (https://github.com/tseemann/mlst) and the scheme of the University of Warwick [6]. Finally, the tree was annotated using Itol [7].

To analyse the genetic environment of the *mhp* gene cluster among B2 strains, we extracted nucleotide sequences ranging from *prpR* to *hemB* genes among all B2 genomes. In the absence of *prpR*, we extracted sequences up to 16,771 bp upstream of *mhpR* to obtain an equivalent distance. Then, we clustered sequences using CD-HIT [8] with identity threshold of 99% and coverage of 95% and kept one represent for each cluster. From the 16 sequences obtained, we construct a physical map using clinker [9] using standard parameters. We annotated the map with the sequence type of the strains and, in the case of ST131, assigned the strains to  t clades A (O16:H5/*fimH41*, B (O25B:H4/*fimH161*, O25B:H4/*fimH22*) or C (O25B:H4/*fimH41*).

*Within-clone diversity*

We assessed the within-clone diversity as the divergence created by the mutations accumulated by each clone during its residence in ED. Conversely to genetic diversity across clones, we were here specifically interested in the differences observed between isolates of the same clone. Therefore, we considered here the clones sampled more than once (two to seven times, for a total of 16 samples considered).

Among those clones, F ST59/815 O1:H7 *fimH*34 presented two distinct haplotypes, both already present in the first sample and differing from each other by six mutations. Both haplotypes were also found in later isolates (the first one 92, 134, 141 and 155 days later; the second one 30 and 489 days later), so that they coexisted in the gut. We performed a clustering analysis on the profiles of all isolates from this clone in order to assign them to one of the two groups, and we assessed the number of differences within each group, compared with the difference between them. The between-cluster variance represented 67.9% of the total, and 1.6 and 1.1 mutations respectively separated two isolates from the same cluster on average, while 7.4 mutations separated two clones belonging to different clusters. Moreover, the six mutations identified in the first sample never appeared in the first group, but always in the second. For the following analyses, we therefore considered these two haplotypes separately.

First, we estimated $v$ the average observed mutation rate of these clones in ED, i.e. excluding highly deleterious mutations that would not be observed. To do so, we computed the average number of mutations separating two isolates of the same sample $i$ (noted $\Pi_{ii}$) and of two separate samples $i$ and $j$ (noted $\Pi_{ij}$). The number of observed mutations accumulated per site over $T$ days separating the two samples ($T = j - i$) could be estimated as $vT = \Pi_{ij} - \Pi_{ii}$ [10]. We computed $vT$ for the 12 samples that were not the first for a given clone (58 isolates), and a linear regression over time was performed to estimate the observed mutation rate per site per day.

Second, we simulated the expected divergence between isolates of the same clone over time under neutrality, using a Wright-Fisher model. Starting from two identical genotypes at time $t = 0$ ($\Pi_{00} = 0$), the average number of mutations separating two isolates is expected to increase towards the mutation-drift equilibrium, depending only on the mutation rate per site per generation (noted $\mu$) and the effective population size (noted $N_e$) [10]. To assess those transient dynamics and compare them to our data, we performed these simulations over 1065 days (the longest time between two samples of the same clone in ED) for various population sizes $N_e$, ranging from 100 to 10000.
